# Caffeine but not acetaminophen increases 4-km cycling time-trial performance

**DOI:** 10.1101/567313

**Authors:** Fabiano Tomazini, Ana Carla S. Mariano, Victor A. Andrade-Souza, Viviane C. Sebben, Carlos A. B. de Maria, Daniel B. Coelho, Romulo Bertuzzi, Marcos D. Silva-Cavalcante, Adriano E. Lima-Silva

**Author notes:** Corresponding author (AEL-S).

## Abstract

Acetaminophen has been combined with caffeine for therapeutic purpose, but the effect of co-ingestion of acetaminophen and caffeine on exercise performance has not been investigated. The aim of this study was to determine the effect of isolated and combined ingestion of caffeine and acetaminophen on performance during a 4-km cycling time-trial. In a double-blind, crossover design, eleven men, accustomed to cycling recreationally, completed a 4-km cycling time-trial one hour after the ingestion of cellulose (PLA), acetaminophen (20 mg·kg^−1^ body mass, ACT), caffeine (5 mg·kg^−1^ body mass, CAF) or combined acetaminophen and caffeine (20 and 5 mg·kg^−1^ body mass, respectively, ACTCAF). The perception of pain and rating of perceived exertion were recorded every 1-km, and electromyography and oxygen uptake were continually recorded and averaged each 1-km. Plasma lactate concentration was measured before and immediately after the trial. The time and mean power during the 4-km cycling time-trial was significantly improved (*P* < 0.05) in CAF (407.9 ± 24.5 s, 241.4 ± 16.1 W) compared to PLA (416.1 ± 34.1 s, 234.1 ± 19.2 W) and ACT (416.2 ± 26.6 s, 235.8 ± 19.7 W). However, there was no difference between ACTCAF (411.6 ± 27.7 s, 238.7 ± 18.7 W) and the other conditions (*P* > 0.05). The perception of pain, rating of perceived exertion, electromyography, oxygen uptake, and plasma lactate were similar across the conditions (*P* > 0.05). In conclusion, caffeine but not acetaminophen increases power output ultimately increasing performance during a 4-km cycling time-trial.

## Introduction

During a self-paced, high-intensity cycling time-trial (e.g., 4-km cycling TT), the exercise intensity must be strictly regulated to avoid an exacerbated early accumulation of metabolites that can lead to fatigue [1–3]. A disturbance in the intramuscular metabolic milieu (i.e., accumulation of the H^+^, ADP, AMP and Pi) at the beginning of a high-intensity cycling TT, provoked by increased muscle recruitment (as inferred from electromyography signals, EMG) ultimately increasing power output (PO) [2,3], will activate peripheral sensory nerve terminals [4,5]. These increased afferent signals might result in increased perception of pain [6], leading to a reduction in PO [2,3].

Acetaminophen (commonly known as paracetamol) has recently been introduced as a potential pharmacological agent to increase exercise performance due to its analgesic proprieties [7–9]. The mechanism by which acetaminophen reliefs pain feelings in humans is not fully known, but it has been attributed to the inhibition of the cyclooxygenase enzymes [10–13], potentiation of descending serotoninergic pathways [14,15], and modulation of opioid and cannabinoid CB1 receptors [11,16]. Although there is not a consensus [17,18], several studies have reported improved performance during high-intensity exercises after a single clinical dose (1-1.5 g or 20 mg·kg^−1^) of acetaminophen [7–9,19].

It is interesting to note that acetaminophen has been combined with caffeine for therapeutic purpose [20,21], but the effect of co-ingestion of acetaminophen and caffeine on exercise performance has not been investigated. The precise mechanism by which caffeine assists acetaminophen in reduction of pain is not fully known, but caffeine might act as an adenosine antagonist at the A2 adenosine receptors of the peripheral sensorial nerves, blocking adenosine-induced pain transmission [22]. The analgesic effect of caffeine may also involve inhibition of presynaptic adenosine receptors on cholinergic nerve terminals in supraspinal sites [23] and increased release of β-endorphin [24]. Although caffeine has other central and peripheral mechanisms of action [25–28] potentially more prominent to exercise performance than analgesia, it would be interesting to explore whether a combination of acetaminophen with caffeine would improve exercise performance to a larger extent than when they are ingested in isolation.

Therefore, the aim of the present study was to investigate the effect of isolated and combined ingestion of acetaminophen and caffeine on performance during a 4-km cycling TT. We hypothesized that exercise performance might improve to a larger extent when acetaminophen and caffeine are combined, when compared to acetaminophen or caffeine alone.

## Materials and Methods

### Participants

Eleven men (age: 24.5 ± 6.9 years, body mass: 74.8 ± 6.5 kg, height: 176.4 ± 7.7 cm, V̇O_2_max 39.0 ± 6.5 ml·kg^−1^.min^−1^), who were accustomed to cycling recreationally, were recruited. Participants were free of acute/chronic pain and were not using acetaminophen or any other analgesic medications. The study was conducted in accordance with ethical standards principles expressed in the Declaration of Helsinki. Participants signed a consent form agreeing to participate in the study, which was approved (written, CAAE: 44457215.20000.5208; Approval: 1.097.620) by the Human Ethics Committee of the Federal University of Pernambuco.

### Experimental Design

Participants visited the laboratory on seven different occasions. At the first visit, participants answered an acetaminophen risk assessment questionnaire [29] and a caffeine consumption questionnaire [30]. No participant presented any risk associated with acetaminophen ingestion. The participant’s habitual consumption of caffeine was 93.3 ± 118.1 mg·day^−1^. Anthropometric measurements and an incremental exercise test were then performed on a road bike (Giant^®^, Aluxx 6061, Taichung, Taiwan) coupled to a pre-calibrated CompuTrainer (CompuTrainer^®^ Pro, RacerMate^®^, Seattle, WA, USA). The seat height was recorded and reproduced for all subsequent sessions. On the second and third visits, at least 72-h apart, participants performed two 4-km cycling TT in each visit for familiarization (separated by 30 min); therefore, participants had the opportunity to practice the 4-km cycling TT four times. The coefficient of variation for exercise time between the two “fresh” trials (first trial of visit 1 and 3) was 1.7 ± 1.3%.

Experimental trials were performed from the fourth to the seventh visits. One hour before the trial, participants ingested gelatin capsule containing either cellulose (PLA), or acetaminophen (20 mg·kg^−1^ body mass, ACT), or caffeine (5 mg·kg^−1^ body mass, CAF), or acetaminophen combined with caffeine (20 and 5 mg·kg^−1^ body mass, respectively, ACTCAF). The substances were encapsulated in capsules of the same colour and shape (Pharmapele^®^, Recife, PE, Brazil). Experimental trials were performed in a randomized, counterbalanced and double-blind manner. A 7-day period between the four experimental trials was adopted for washout. Participants were instructed to cover the 4-km cycling TT as fast as possible during the trials and received visual feedback for distance completed, but not for exercise time, power output, pedal frequency or physiological parameters. Participants were asked to refrain from consuming caffeine-containing substances and from taking any analgesic medications as well as to restrain from doing exercise during the 24h before each experimental trial. Participants were also asked to complete a 24-h food recall before the first experimental trial and to replicate it in the following trials. Participants were inquired before trial whether they had adhered to dietary intake and abstaining from caffeine and analgesic medications. All tests were performed at the same time of day to avoid the circadian variation on TT performance [31].

### Incremental test

After a 5-min warm-up at 50 W, PO was increased 25 W every minute until exhaustion. Participants were required to maintain cadence at 70 rpm throughout the test. Oxygen consumption (V̇O_2_), dioxide carbon production (V̇CO_2_), and ventilation (V̇E) were measured V VV breath-by-breath during the test through an automatic gas exchange analyser (Cortex, Metalyzer^®^ 3B, Leipzig, Germany). The calibration of the equipment was performed according to the manufacturer’s recommendations using gases of known concentration (12% O_2_ and 5% CO_2_) and a 3-L syringe for the volume. Heart rate (HR) was measured via an HR transmitter coupled to the gas analyser (Polar^®^, T 31/34, Kempele, Finland).

The V̇O_2_max was determined as the highest 20-s average V̇O_2_ during the test. Aerobic peak power was determined as the PO achieved during the last completed stage. When the participants were not able to maintain the PO during an entire stage (i.e., <1 min), the aerobic peak power was calculated using the fractional time completed in the last stage completed, multiplied by the increment rate [32].

### Experimental trials

Participants arrived at the laboratory, after a 3-h fasting, and a double Ag/AgCl electrode was attached over the *vastus lateralis* belly for further EMG record [33]. To further normalize the EMG signal during the trials, participants performed three maximal voluntary contractions (MVC) with a 1-min passive rest between the contractions on a custom-made bench. The hip and knees angles were set at 120° and 90°, respectively. A non-compliant cuff attached to a calibrated linear strain gauge was fixed to the right ankle just superior to the malleoli for force measurement. The EMG and force signals were acquired with a sample rate of 2,000 Hz (EMG System, EMG 830 C, São Jose dos Campos, SP, Brazil). The EMG signal during the MVC was further used to normalize the EMG signal during the 4-km cycling TT.

After MVCs recording, participants ingested capsules containing PLA, ACT, CAF, or ACTCAF and rested for 45 min (Fig 1). Then, participants were questioned about which substances they thought they had ingested, and then performed a 5-min warm-up at 50% of aerobic peak power, followed by a 5-min rest and a 4-km cycling TT. Participants were instructed to cover the 4-km cycling TT as fast as possible. The trials were performed in an ambient temperature of 21 ± 1 °C and a relativity humidity of 38 ± 5 %.

**Fig 1.**
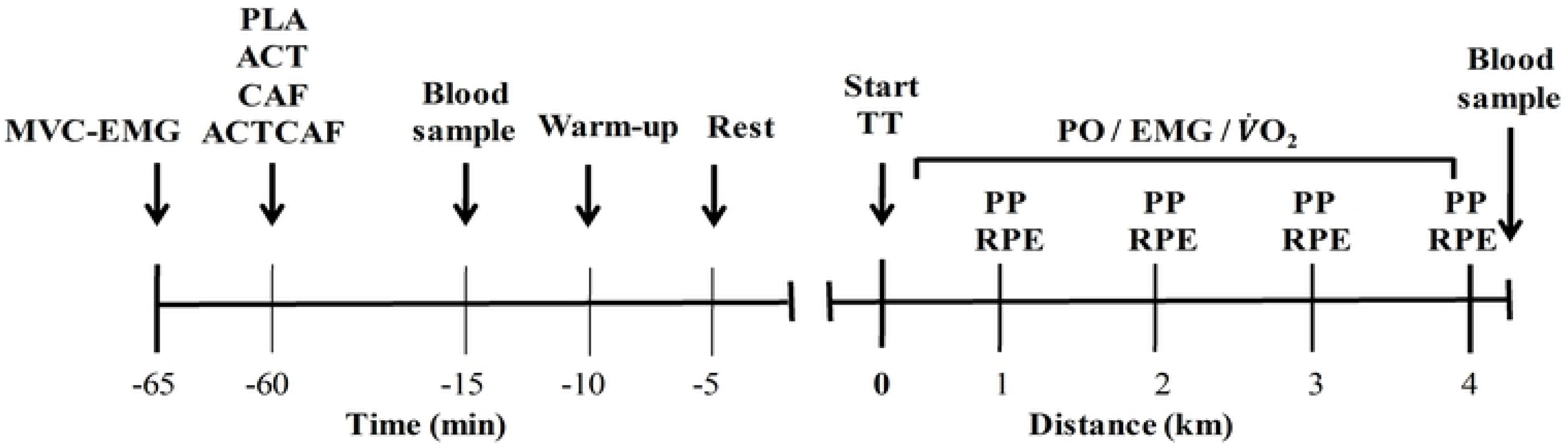
Timeline of the experimental trial. Maximal voluntary contraction (MVC), eletromyographicic activity (EMG), placebo (PLA), acetaminophen (ACT), caffeine (CAF) or acetaminophen combined with caffeine (ACTCAF), time trial (TT), power output (PO), oxygen uptake (V̇O_2_), perception of pain (PP), and rating of perceived exertion (RPE).

The PO was recorded every second during the entire trial (RacerMate Software, version 4.0.2, Seattle, WA, USA). The V̇O_2_ and EMG were continually recorded. The RPE and pain perception were recorded every 1-km using the 15-point Borg’s scale [34] and the 12-point pain intensity scale [35], respectively. Capillary blood samples (50 μL) were collected from the earlobe in a sodium-heparinized capillary tube before and immediately after the trial.

### Data analyses

The raw EMG signal was filtered with a fourth-order Butterworth band-pass filter with cut-off frequencies set at 10 and 400 Hz. Signal was full-wave rectified and root mean square (RMS) calculated for a 500-ms window around peak torque during MVC. The RMS of each burst during the 4-km cycling TT was also calculated, averaged for every 1-km epoch and normalized by RMS of MVC. The PO and V̇O_2_ data were also averaged every 1-km interval. Blood samples were centrifuged at 4000 rpm (4°C) during 15 minutes for plasma separation, and plasma lactate concentration ([La]) was determined with commercial kits (Labtest, Lagoa Santa, MG, Brazil) with the reaction reading in a spectrophotometer (Thermo Scientific, Genesis 10S UV-Vis, Waltham, MA, USA).

### Statistical analyses

The assumption of normality was checked with the Shapiro-Wilk test. The performance time was compared between the conditions via ANOVA of repeated-measures. The dependent variables measured during the TT were compared between conditions using a 2-way ANOVA of repeated-measures (condition x distance). Ducan’s *post hoc* test was conducted when a significant main effect or interaction was detected. The level of significance adopted was α < 0.05. Data were analysed using the statistical package Statistica (StaSoft^®^, Inc. version 10, Tulsa, OK, USA).

## Results

### Blinding

From 44 trials, participants guessed 16, 14, 7 and 5% the ingestion of PLA, ACT, CAF and ACTCAF, respectively, indicating a success in the blinding process.

### Exercise Performance

There was a significant effect of substance ingestion on exercise performance (*P* = 0.029, Fig 2). The time to complete the 4-km cycling TT was significantly faster in CAF (407.9 ± 24.5 s) than in PLA (416.1 ± 34.1 s, *P* = 0.015) and ACT (416.2 ± 26.6 s, *P* = 0.016), but there was no significant difference between PLA and ACT (*P* = 0.976). There was also no significant difference between ACTCAF (411.6 ± 27.7 s) and the other three conditions (*P* > 0.05).

**Fig 2.**
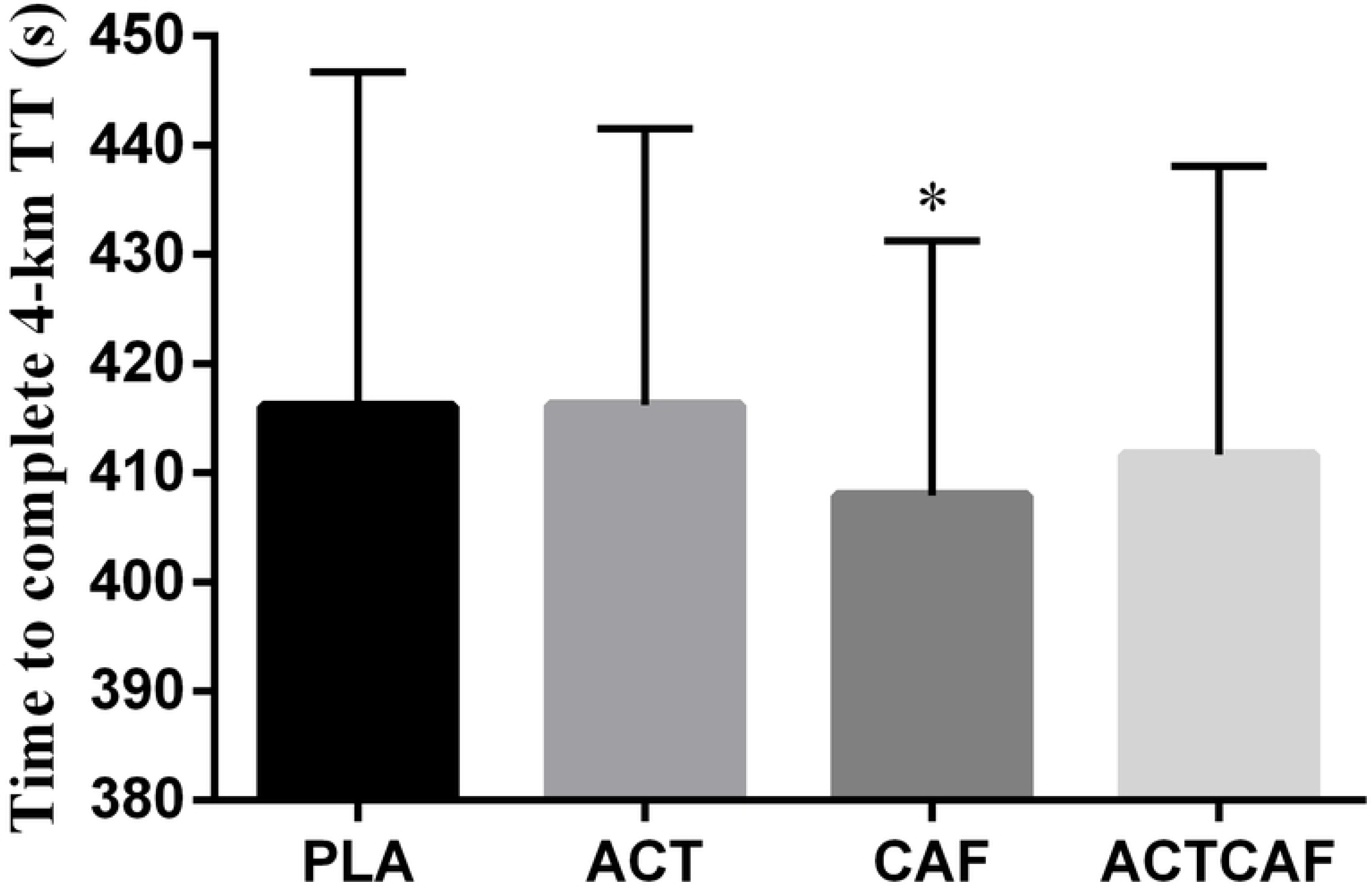
Performance time during a 4-km cycling TT after placebo (PLA), acetaminophen (ACT), caffeine (CAF) and acetaminophen combined with caffeine (ACTCAF) ingestion. Data are shown as mean ± SD. *Significantly faster than PLA and ACT (P < 0.05).

### Power output distribution, neuromuscular recruitment, and metabolic and perceptual responses

There was a main effect of condition (*P* = 0.022) and distance (*P* = 0.008), but not condition x distance interaction (*P* = 0.795) for PO (Fig 3A). The PO was higher throughout the trial in CAF (241.4 ± 16.1 W) compared to PLA (234.1 ± 19.2 W, *P* = 0.007) and ACT (235.8 ± 19.7 W, *P* = 0.031), without any other differences (all *P* > 0.05). The PO was higher at the beginning and end of the-trial, compared to the middle of the trial (*P* = 0.012 and *P* = 0.049, respectively). There was only a main effect of distance for RMS (*P* = 0.027, Fig 3B), V̇O_2_ (*P* = 0.001, Fig 3C), RPE (*P* = 0.001, Fig 3D), and pain perception (*P* = 0.001, Fig 3E), without any condition or condition x distance interaction (*P* > 0.05). Plasma lactate increased after the trial (*P* = 0.001), but there was no condition (*P* = 0.322) or condition x distance interaction (*P* = 0.336).

**Fig 3.**
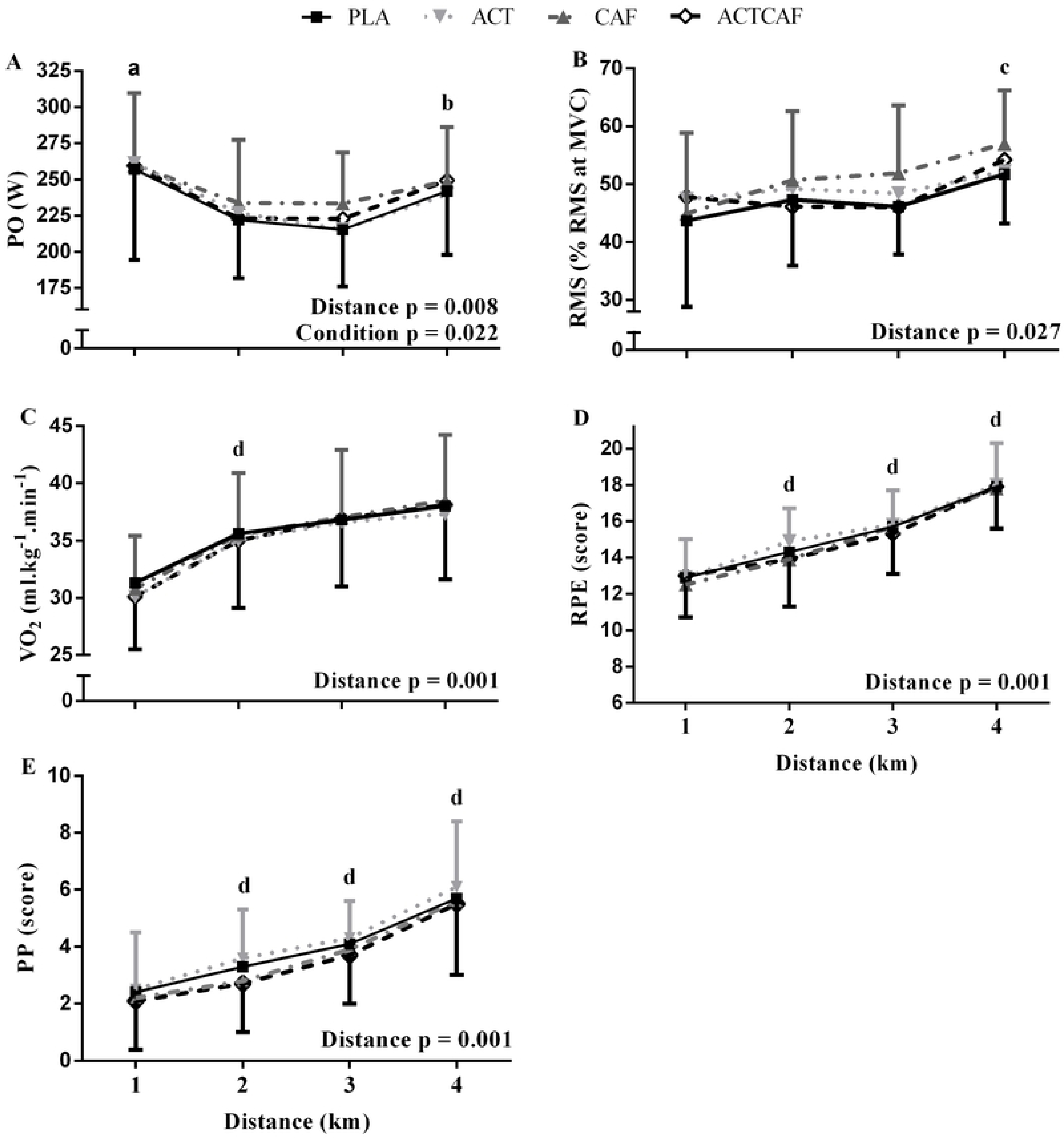
Physiological and neuromuscular parameters during a 4-km cycling time-trial after placebo (PLA), acetaminophen (ACT), caffeine (CAF) or acetaminophen combined with caffeine (ACTCAF) ingestion. (A) Power output. (B) Electromyography. (C) Oxygen uptake. (D) Rating of perceived exertion. (E) Pain perception. Data are shown as mean ± SD. (A) PO was higher in CAF than in PLA and ACT throughout the trial (main effect of condition). ^a^Significantly higher than in 2 and 3 km (*P* < 0.05), ^b^Significantly higher than in 3-km (*P* < 0.05). (B) ^c^Significantly higher than all previous points (*P* < 0.05). (C, D and E) ^d^Significantly higher than previous one (*P* < 0.05).

## Discussion

The present study is the first comparing the effect of isolated and combined caffeine and acetaminophen ingestion on exercise performance. Our findings indicated that the isolated ingestion of caffeine increases power output throughout a 4-km cycling TT, ultimately improving exercise performance, when compared to placebo and acetaminophen alone. However, this effect is lacking when caffeine is combined with acetaminophen. Acetaminophen alone had no effect on exercise performance.

### Acetaminophen alone

In the present study, the ingestion of acetaminophen alone (20 mg·kg^−1^) did not increase 4-km cycling TT performance. Based on previous findings [7,9], the acetaminophen ingestion was expected to improve exercise performance. This non-improvement in exercise performance after acetaminophen ingestion in the present study conflicts with previous observations that acetaminophen ingestion (1.5 g) improves exercise performance during a 16.1-km cycling TT [9], increases total work done during a 3-min all-out test [36], increases time to task failure at 70% V̇O_2_max (8), raises the peak or mean power output during repeated sprints [7,19], and increases mean torque during a 60 x 3-s MVC (2-s passive recovery period) protocol [37]. On the other hand, other studies failed to find an effect of acetaminophen ingestion on time to task failure at 70% V̇O_2_max [17] or repeated sprint performance [18]. The reason for these contradictory results is not clear, but it may be related to different exercise protocols and tasks measuring exercise performance. Our findings add to these showing that acetaminophen ingestion (20 mg·kg^−1^) did not improve exercise performance during a self-paced, high-intensity exercise such as a 4-km cycling TT. In view of these conflicting results, it could be argued that the ergogenicity of the acetaminophen might be task-dependent. It is recommended therefore that acetaminophen should be tested in TT of different lengths.

### Caffeine alone

In the present study, caffeine ingestion (5 mg·kg^−1^) improved exercise performance by ∼2%, when compared to both placebo and acetaminophen. These findings are consistent with several other studies showing a positive effect of caffeine compared to placebo (2-2.4%) on performance in a 4-km cycling TT [26,38,39]. However, the comparison between the ergogenicity of caffeine and acetaminophen had not been investigated; therefore, our findings suggest that caffeine has a greater ergogenic potential than acetaminophen, at least during a high-intensity cycling TT as used in the present study.

Different mechanisms have been proposed to explain the ergogenicity of caffeine [27,38,39]. One of them is that caffeine might have analgesic proprieties due to its peripheral effect on A_2_ adenosine receptors [22] and/or on central β-endorphin release [40]. In the present study, caffeine alone led to greater mean cycling power output, with the same level of perceived pain and effort. This finding agrees with previous studies showing an increased power output with caffeine during high-intensity exercise, without changes in muscle pain [41,42] or RPE [26,38,43].

It should be highlighted that caffeine has other physiological effects not directly related to pain relief, which might be more important to improve exercise performance. In the present study, an increased power output with caffeine supplementation was not accompanied by an increased muscle recruitment, as inferred from results of EMG. Caffeine is expected to have a central effect due its action as an antagonist of the A1 adenosine receptors in the central nervous system [42]. It is predictable therefore that caffeine ought to increase muscle recruitment. However, data from the literature are contradictory, with some studies showing an increased muscle recruitment with caffeine during a 4-km cycling TT [26] and lower reduction in voluntary activation of the *vastus lateralis* during intermittent isometric knee extension contractions [44], while others failed to find any effect of caffeine on motor unit recruitment [38,45,46]. These contradictory findings may be because caffeine can also influence spinal excitability and the muscle contractile apparatus. Caffeine increases spinal excitability [47], which would demand less central neural drive to maintain muscle activation. Caffeine in a physiological concentration in plasma (∼40 µmol·L^−1^) also improves calcium release from the sarcoplasmic reticulum, which increases muscle force for a given level of muscle recruitment [48–50].

### Combined acetaminophen and caffeine ingestion

Clinical trials have demonstrated that the combination of acetaminophen (500-2000 mg) and caffeine (65-260 mg) reduces the time of onset of analgesic action and increase analgesic effectiveness, compared to acetaminophen alone [20,21]. We found a non-significant effect of the combination of acetaminophen and caffeine on exercise performance, when compared to placebo or to isolated ingestion of ACT or CAF. However, although not significant, time to cover the 4-km was slightly faster in ACTCAF compared to PLA (∼1%), but slightly slower compared to CAF (∼1%). The mechanism by which acetaminophen would negatively influence caffeine-induced improvement in exercise performance cannot be provided from our data. Further studies should test different combinations of caffeine and acetaminophen doses to determine if there is a dose combination most able to improve exercise performance. At the time of our study, however, the combination of acetaminophen with caffeine is not recommended.

### Limitations

The present study has some limitations that should be mentioned. We recruited participants who did cycling recreationally, whose day-by-day variation may be greater than in trained cyclists. However, participants performed four familiarization trials, which helped to reduce the potential effect of learning. This assumption is supported by the low coefficient of variation between the two “fresh” familiarization trials (∼1.7%), which is very close to that reported for trained cyclists (∼1.9%) [51].

## Conclusion

In conclusion, acute caffeine ingestion but not acetaminophen increases cycling power output, improving performance of a 4-km cycling TT. The acetaminophen seems, however, to negatively affect the ergogenicity of caffeine. Therefore, the combined use of caffeine and acetaminophen should be avoided.

## Acknowledgments

The authors thank the participation of all volunteers of the study. The authors also thank the technical assistance of Thaysa Ghiarone, Kleiton Silva, Guilherme Ferreira and Leandro Felippe. The English text of this paper has been revised by Sidney Pratt, Canadian, MAT (The Johns Hopkins University), RSAdip - TESL (Cambridge University).

